# Extending the spacing between the Shine-Dalgarno sequence and P-site codon reduces the rate of mRNA translocation

**DOI:** 10.1101/2020.04.16.045807

**Authors:** Hironao Wakabayashi, Chandani Warnasooriya, Dmitri N. Ermolenko

**Author notes:** To whom correspondence may be addressed: Dmitri N. Ermolenko.

## Abstract

By forming basepairing interactions with the 3’ end of 16S rRNA, mRNA Shine-Dalgarno (SD) sequences positioned upstream of Open Reading Frames (ORFs) facilitate translation initiation. During the elongation phase of protein synthesis, intragenic SD-like sequences stimulate ribosome frameshifting and may also slow down ribosome movement along mRNA. Here, we show that the length of the spacer between the SD sequence and P-site codon strongly affects the rate of ribosome translocation. Increasing the spacer length beyond six nucleotides destabilizes mRNA-ribosome interactions and results in a 5-10 fold reduction of the translocation rate. These observations suggest that during translation, the spacer between the SD sequence and P-site codon undergoes structural rearrangements, which slow down mRNA translocation and promote mRNA frameshifting.

## Introduction

The <u>S</u>hine-<u>D</u>algarno (SD) sequence plays several important roles in regulation of protein synthesis in bacteria. 3-to-9 nucleotide-long SD sequences (UAAGGAGGU) occur upstream of the start codon of most bacterial ORFs and increase efficiency of translation initiation through basepairing interactions with the anti-SD (aSD) sequence at the 3’ end of 16S rRNA [1, 2].

SD-like sequences, such as Arg (AGGAGG) or Gly (GGAGGU) codon pairs, also occur inside <u>o</u>pen reading frames (ORFs) where they may regulate translation elongation. Ribosome profiling studies [3, 4] and *in vitro* single-molecule experiments [5, 6] suggested that SD-like sequences present within ORFs of mRNA induce ribosome pausing. Specific internal SD-like sequences within ORFs are conserved between divergent bacterial species [3] indicating that SD-induced pauses may be functionally important. SD-induced ribosome pausing was suggested to prevent the formation of anti-termination RNA stem-loops and, thus, enable proper transcription termination [3]. In addition, clustering of SD-like sequences in mRNA segments coding for loops in protein structures were hypothesized to modulate co-translational protein folding [3]. However, more recent ribosome profiling [7] and biochemical studies [8] showed that intragenic SD-like sequence have little or no effect on the rate of ribosome translocation along mRNA.

While it remains unclear whether SD-like sequence induce ribosome pausing, involvement of SD sequences in <u>p</u>rogrammed ribosome frameshifting (PRF) has been well documented [9-11]. For example, a SD sequence induces +1 PRF, which produces release factor 2 (RF2) in *E. coli*. SD sequences also stimulate -1 PRF events that produce γ subunit of DNA polymerase III and cytidine deaminase [11-14].

Elucidation of the effect of SD sequences on elongation may be complicated by the fact that the strength of the SD-aSD interaction differs from mRNA to mRNA as the number of SD-aSD basepairs varies from 3 to 9 nucleotides. Furthermore, the spacer between the SD sequence and P-site codon was shown to have a strong effect on the efficiency of both translation initiation and SD-stimulated frameshifting in bacteria [9, 10, 15, 16]. For example, SD sequences positioned less than 4 nucleotides upstream of the P site codon destabilize P-site binding of tRNA and promote +1 PRF [9, 16]. The most optimal spacing between SD and the A of AUG start codon for the promotion of translation initiation is 4-9 nucleotides [15]. By contrast, the most optimal spacing between the P site codon and SD sequences for stimulation of -1 PRF in *dnaX* mRNA is 10-14 nucleotides [10]. The stimulatory effect of the SD sequence on -1 PRF in *dnaX* mRNA disappears when the spacer between SD and the slippery sequence is increased from 10 to 16 nucleotides or decreased from 10 to 7 nucleotides [10]. It is not clear why the length of the spacer between the SD sequence and P-site codon optimal for initiation is significantly different from the spacing optimal for the -1 PRF.

Ribosome profiling data [4] and indirect biochemical experiments [17] suggest that SD-aSD interactions can be preserved for a number of rounds of ribosome translocation that possibly results in looping or inchworm movement of the spacer sequence between the SD and P-site codon inside the ribosome. At the point of SD-aSD hybrid dissociation, the hybrid may create tension that pulls the ribosome back and induces ribosome pausing and frameshifting [11]. It is likely that the effect of the SD sequence on translocation depends on the number of SD-aSD basepairs and the spacing between the SD and translocating codons in mRNA. Thus, variations in these parameters may underlie the discrepancies between different studies of the SD inhibitory effect on ribosome translocation.

Here we (i) determine to what extent the SD-anti-SD interaction slows down ribosome translocation and (ii) elucidate the dependence of translocation rate on stability of the SD-anti-SD duplex and the spacing between the SD sequence and P-site codon. Our results are consistent with the idea that SD-aSD interactions are retained for several rounds of ribosome translocation along mRNA. We also find that the effect of SD-aSD interactions on translocation is highly variable and depends on the spacing between the SD sequence and P-site codon. Our results reconcile previous contradictory reports regarding the role of SD-like sequences in regulating the rate of translation elongation and provide new insights into mechanisms of ribosome pausing and frameshifting.

## Results

### Lengthening the spacer between the SD and P-site codon destabilizes mRNA-ribosome interactions

To test how the length of the spacer between the SD sequence and P site codon affect stability of ribosome-mRNA interactions and ribosome translocation, we made eight model mRNAs based on the derivative of the phage T4 gene *32* mRNA named m301 [18, 19]. In these eight mRNA, we varied spacing between a AAGGA SD sequence and a UAC (Tyr) codon from 6 to 21 nucleotides. We first examined an initiation-like complex containing a tRNA in the P site. We assembled ribosomal complexes with each mRNA variant and a deacylated tRNA^Tyr^ and measured stability of ribosome-mRNA interactions using toeprinting. In toeprinting assay, the position of the ribosome along mRNA is mapped by inhibition of reverse transcriptase that extends DNA primer annealed downstream of the ribosome. The ribosome containing P-site tRNA produces a toeprint 16 nucleotides downstream from the first nucleotide of the P site codon. The presence of SD-aSD interactions stabilizes the mRNA-ribosome complex and substantially increases intensity of the toeprint [16, 18]. Comparison of toeprint intensities between mRNAs with different SD-UAC spacing indicated that stability of mRNA-ribosome complex progressively decreased as spacing between the SD sequence and P-site codon was extended from 6 to 11 nucleotides (Fig. 1). Toeprints became barely detectable when spacing between the SD and P-site codon was extended beyond 11 nucleotides. These results suggest that elongating spacing between the SD sequence and P site codon beyond the length which was shown to be optimal for stimulating translation initiation, destabilizes mRNA-ribosome interactions.

**Figure 1.**
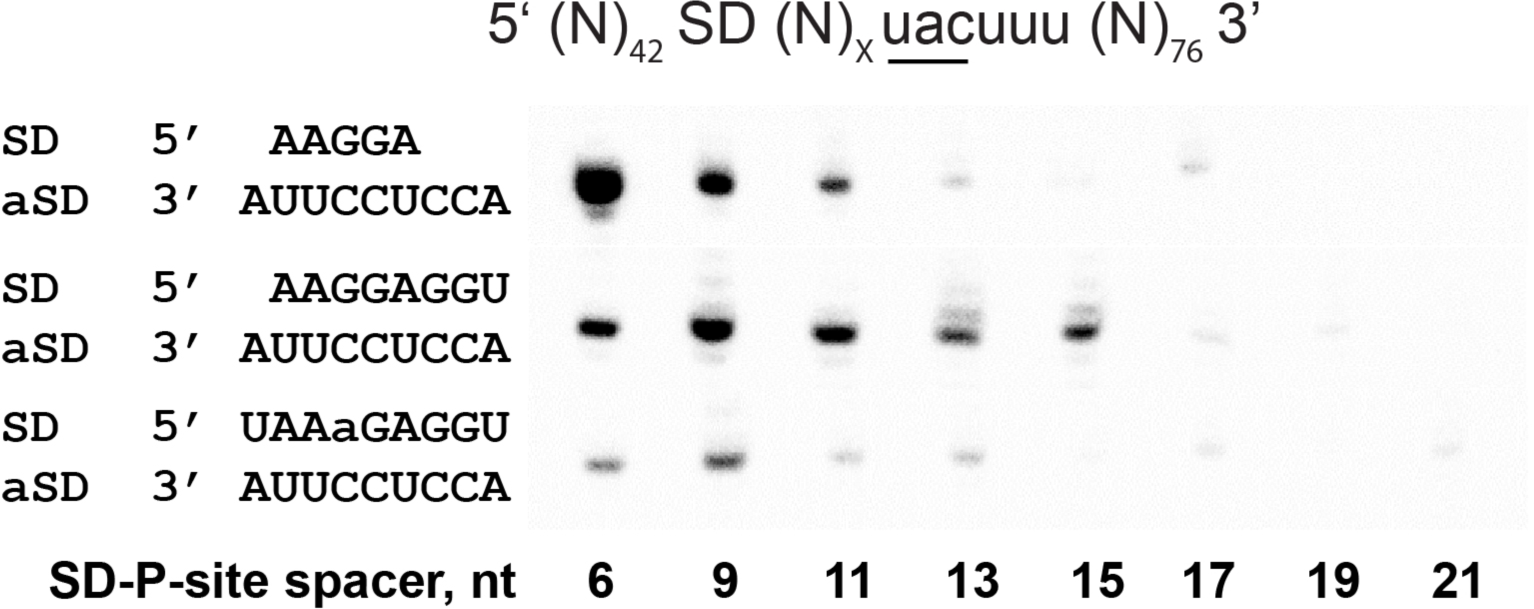
Extending spacing between the SD sequence and the P site codon destabilizes mRNA-ribosome interactions. Toeprints were produced by incubating the 70S ribosome with an mRNA and tRNA^Tyr^, which basepairs with a unique UAC codon downstream of the SD. Each mRNA contained either AAGGA, AAGGAGGU or UAAaGAGGU SD sequence as indicated. The spacer between the SD and UAC codon varied between 6 and 21 nucleotides in length as indicated.

We next asked how the spacer length affects mRNA-ribosome interactions in an elongation complex, which contains two tRNAs. Binding of an A-site tRNA stabilized mRNA-ribosomes interactions. In contrast to ribosome complexes containing a single P-site tRNA (Fig. 1), pre-translocation elongation complexes containing both A- and P-site tRNAs produced detectable toeprints even when spacing between the SD sequence and UAC codon was as long as 21 nucleotides (Suppl. Fig. 1). Nevertheless, the dependence on the spacer length followed the same trend as for a single tRNA, initiation-like complex. The stability of mRNA-ribosome complex progressively decreased as the spacing between the SD sequence and P-site codon increased from 6 to 21 nucleotides (Suppl. Fig. 1).

We next tested whether the dependence of ribosome-mRNA interactions on spacing between the SD and P-site codon is altered when the strength of SD-aSD interactions is increased. To that end, we replaced the AAGGA SD sequence in all eight original mRNA variants with AAGGAGGU that is predicted to increase the thermodynamic stability of SD-aSD duplex from -3.6 kcal/mol to -11.2 kcal/mol. In contrast to mRNAs containing AAGGA SD sequence, extending spacing between AAGGAGGU SD sequence and UAC codon from 6 to 9 nucleotides increased intensity of the toeprint produced by the ribosome containing P-site tRNA^Tyr^. Further extension of the spacer between the SD and UAC codon gradually decreased the toeprint intensity, yet the toeprint remain readily detectable until the spacing was extended to 17 nucleotides. Thus, AAGGAGGU SD sequence increased stability of the ribosome-mRNA complex relative to AAGGA SD sequence. In addition, SD sequences AAGGA and AAGGAGGU differ in optimal spacing between the SD and P-site codon, at which mRNA-ribosome interaction are most stable (6 and 9 nucleotides, respectively).

We further examined whether these differences in optimal SD-P-site spacing at which mRNA-ribosome interaction are most stable, are due to differences in strength of SD-aSD interactions or due to distinct alignments of SD sequences with the aSD sequence of 16S rRNA that could affect SD-P-site spacing. The AAGGA SD sequence anneals to 16S rRNA near the 3’ end of aSD sequence (nt 1537-1541 in *E. coli*) while AAGGAGGU SD sequence spans the entire aSD sequence including its 5’ end (nt 1534-1541 in *E. coli*). To destabilize SD-aSD interactions for the long SD sequence, we introduced one mismatch in SD-aSD duplex by replacing AA<u>G</u>GAGGU SD with UAA<u>a</u>GAGGU SD (lower case letter “a” indicates mismatched adenine) that is predicted to change thermodynamic stability of SD-aSD duplex from -11.2 kcal/mol to -6.4 kcal/mol. A mismatch in SD-aSD duplex decreased intensity of toeprints relative to toeprints observed in the presence of mRNAs containing AA<u>G</u>GAGGU SD, indicating lower stability of mRNA-ribosome complex in ribosomes programmed by mRNAs with UAA<u>a</u>GAGGU SD. However, similar to mRNAs with AA<u>G</u>GAGGU SD, the optimal spacing between UAA<u>a</u>GAGGU SD and P-site codon, at which mRNA-ribosome interactions are most stable, was 9 nucleotides. Hence, optimal SD-P-site spacing at which mRNA-ribosome interaction are most stable is at least partially defined by the alignment of SD sequences relative to aSD sequence.

Taken together, toeprinting experiments indicate that each SD sequence has a specific optimum for spacing between the SD and P-site codon, at which mRNA-ribosome interactions are most stable. Extending the spacer between the SD and P-site codon beyond this optimal length destabilizes mRNA-ribosome interactions. These results suggest that SD-aSD interactions may be maintained for several rounds of translation elongation resulting in inchworm-like movement of the spacer between the SD sequence and P-site codon. However, this rearrangement of the spacer destabilizes mRNA-ribosome interactions.

### Extending the spacing between the SD and P-site codon promotes mRNA back-slippage

Inside ORFs, SD sequences are known to promote PRF. We used our model mRNAs to examine how spacing between the SD and P-site codon affect frame maintenance. It has been previously demonstrated that translocation of deacylated tRNA (but not peptidyl-tRNA) from the A to P site of the ribosome can be accompanied by a repairing of the tRNA anticodon with the upstream synonymous codon, which is positioned closer to the SD sequence (Fig. 2) [18, 20]. This effect has been called mRNA back-slippage. Such mRNA back-slippage is thought to be driven by the SD sequence positioned at a more favorable distance from the upstream codon [18, 20]. The mechanism of this phenomenon, which is observed *in vitro*, is likely similar to the mechanism of SD-driven -1 PRF occurring *in vivo*.

**Figure 2.**
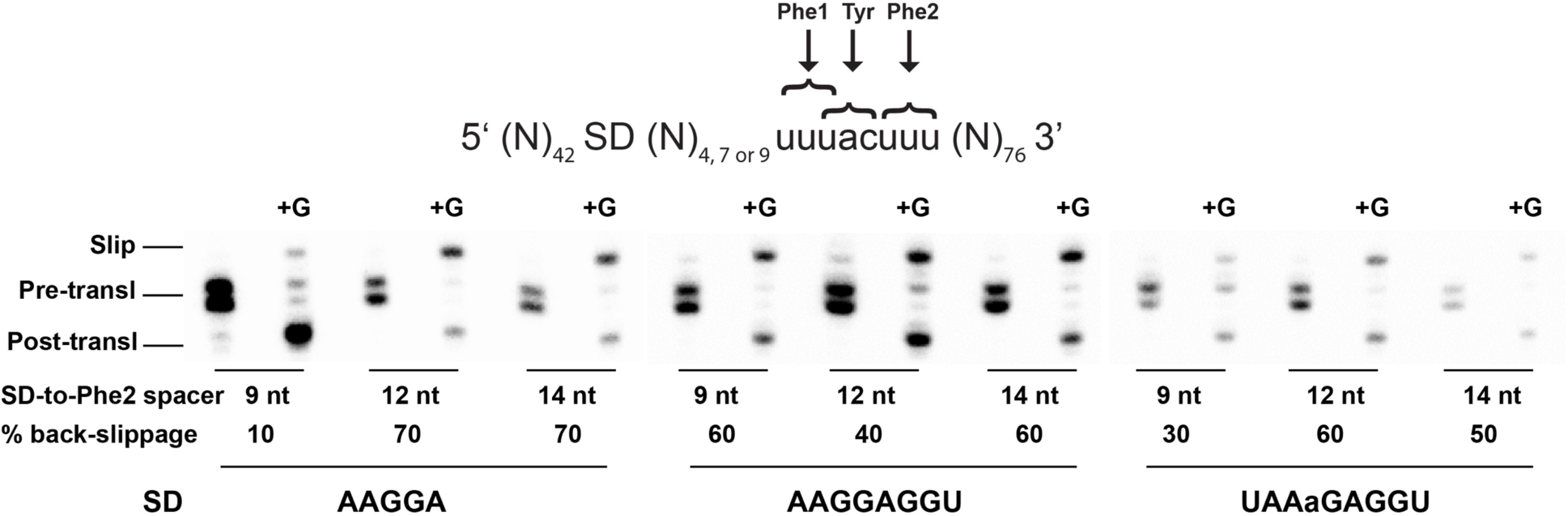
Extending spacing between the SD and P-site codon promotes mRNA back-slippage. A pre-translocation complex (“Pre-transl”) was made by binding deacylated tRNA^Tyr^ to the P site and deacylated tRNA^Phe^ to the A site in the presence of an mRNA. Each mRNA contained either AAGGA, AAGGAGGU or UAAaGAGGU SD sequence as indicated. The spacer between the SD and Phe2 UUU codon was either 9, 12 or 14 nucleotides as indicated. The position of the ribosome along the mRNA was mapped by toeprinting. Pre-translocation complexes were incubated with EF-G and GTP to induce mRNA translocation (+G lanes). “Post-transl” toeprint bands correspond to the product of accurate translocation, in which P-site tRNA^Phe^ is basepaired with Phe2 UUU codon. “Slip” bands correspond to the product of mRNA back-slippage, in which P-site tRNA^Phe^ is basepaired with Phe1 UUU codon. Percent of back-slippage was calculated from the intensity of “slip” band normalized to the sum of intensities of “slip” and “post-transl” bands.

We used our model mRNAs to examine how the spacing between the SD and P-site codon affects frame maintenance. mRNA back-slippage was previously observed in mRNA 301 [19, 21], from which we derived aforementioned model mRNAs. In these model mRNAs, downstream of SD sequence, there are two alternative codons for tRNA^Phe^ flanking the UAC (Tyr) codon (Fig. 2). To test whether efficiency of mRNA back-slippage depends on the spacing between SD sequence and the two Phe codons, toeprinting experiments were performed. Deacylated tRNA^Tyr^ was first bound to the P site of the ribosome programmed with one of our model mRNAs. Next, deacylated tRNA^Phe^ was bound to the A site to basepair with the downstream (Phe2) codon. The formation of pre-translocation complex results in the appearance of the doublet toeprint characteristic of the ribosome containing an A-site tRNA (Fig. 2). Translocation induced by EF-G and GTP resulted in the appearance of a toeprint that corresponds to the post-translocation ribosome containing P-site tRNA^Phe^ basepaired with the downstream Phe codon (“post-transl”, Fig. 2). mRNA back-slippage produced additional toeprint that corresponds to the ribosome containing P-site tRNA^Phe^ basepaired with the upstream (Phe1) codon (“slip”, Fig. 2).

Efficiency of mRNA back-slippage was negligible (≤ 10% of ribosomes) when the spacing between the AAGGA SD and the downstream Phe codon (Phe2) was 9 nucleotides. However, extending spacing between SD and the downstream Phe codon (Phe2) to 12 or 14 nucleotides increased back-slippage efficiency to 70% (Fig. 2). Our toeprinting experiments show that extending the spacing between the SD and the downstream Phe codon (Phe2) destabilizes codon-anticodon interactions and stimulates re-pairing of P-site tRNA^Phe^ with an upstream Phe codon (Phe1), which is positioned five nucleotides closer to the SD sequence.

The stronger AAGGAGGU sequence with the SD-Phe2 spacing of 9 nucleotides has a substantially higher back-slippage efficiency (60%) than that of AAGGA (Fig. 2). Introducing a mismatch into the SD-aSD helix by changing AAGGAGGU to UAAaGAGGU decreased back-slippage efficiency from 60% (with AAGGAGGU SD) to 30% (with UAAaGAGGU SD) (Fig. 2). Similarly to AAGGA, the UAAaGAGGU sequence with the SD-Phe2 spacing extended to 12-14 nucleotides increases efficiency of mRNA back-slippage from 30 to 50-60% (Fig. 2).

Our toeprinting experiments suggest that the mechanism of SD-stimulated -1 PRF in bacteria involves SD-driven destabilization of codon-anticodon interactions, which leads to mRNA back-slippage. This SD-driven destabilization of codon-anticodon interactions depends on spacing between SD and P-site codon. Our toeprinting results are consistent with experiments demonstrating that shortening spacing between P-site codon and the SD sequence from 10 to 7 nucleotides inhibits -1 PRF in *dnaX* mRNA [10].

### Extending the spacing between the SD and P-site codon inhibits ribosome intersubunit rotation coupled to mRNA translocation

We next examined how the strength of SD sequence and spacing between SD and P-site codon affect the rate of ribosome translocation. Ribosome translocation is coupled to cyclic rotational movement between ribosomal subunits [22, 23]. Upon formation of a new peptide bond, ribosomal subunits rotate relative to each other by up to 10° from a nonrotated conformation into a rotated conformation [22, 23]. Binding of EF-G transiently stabilizes the rotated conformation, and catalyzes mRNA translocation during the reverse rotation between subunits [22, 23].

Förster resonance energy transfer (FRET) between fluorophores attached to ribosomal proteins S6 and L9 was previously extensively used to follow intersubunit rotation [24-28]. The reverse intersubunit rotation from rotated to nonrotated conformation increases FRET between fluorophores attached to proteins S6 and L9. Here, we used the FRET assay to examine how SD sequences and SD-P-site spacing affect translocation. To this end, we measured pre-steady-state kinetics of the reverse rotation between subunits of the ribosomes programmed with aforementioned model mRNAs, which differ in SD sequences and spacing between P-site codon and SD.

To assemble pre-translocation ribosomes, deacylated tRNA^Tyr^ was first bound to the P site of the ribosome programmed with a model mRNA. Next, N-acetyl-Phe-tRNA^Phe^ was bound to the A site. Consistent with published reports [18, 20], no mRNA back-slippage was observed when peptidyl-tRNA (N-acetyl-Phe-tRNA^Phe^) translocated from the A to P site upon addition of EF-G and GTP (Suppl. Fig. 1).

When pre-translocation ribosomes were mixed with EF-G and GTP in a stopped-flow fluorometer, we observed a rapid increase in acceptor fluorescence indicating increase in FRET (Fig. 3a). The FRET increase corresponds to the reverse rotation of ribosomal subunits into non-rotated conformation of the ribosome that accompanies mRNA translocation [26]. Kinetic traces were best fitted to a double-exponential function. Bi-phasic kinetics of ribosome translocation was previously observed in multiple studies [26, 29-31]. It is not fully known whether the bi-phasic kinetics of translocation is due to heterogeneity of ribosome population or other factors. The faster rate constant *k*_*1*_, which accounts for 55-70% of the amplitude of fluorescence change, is typically used as a measure of translocation rate.

**Figure 3.**
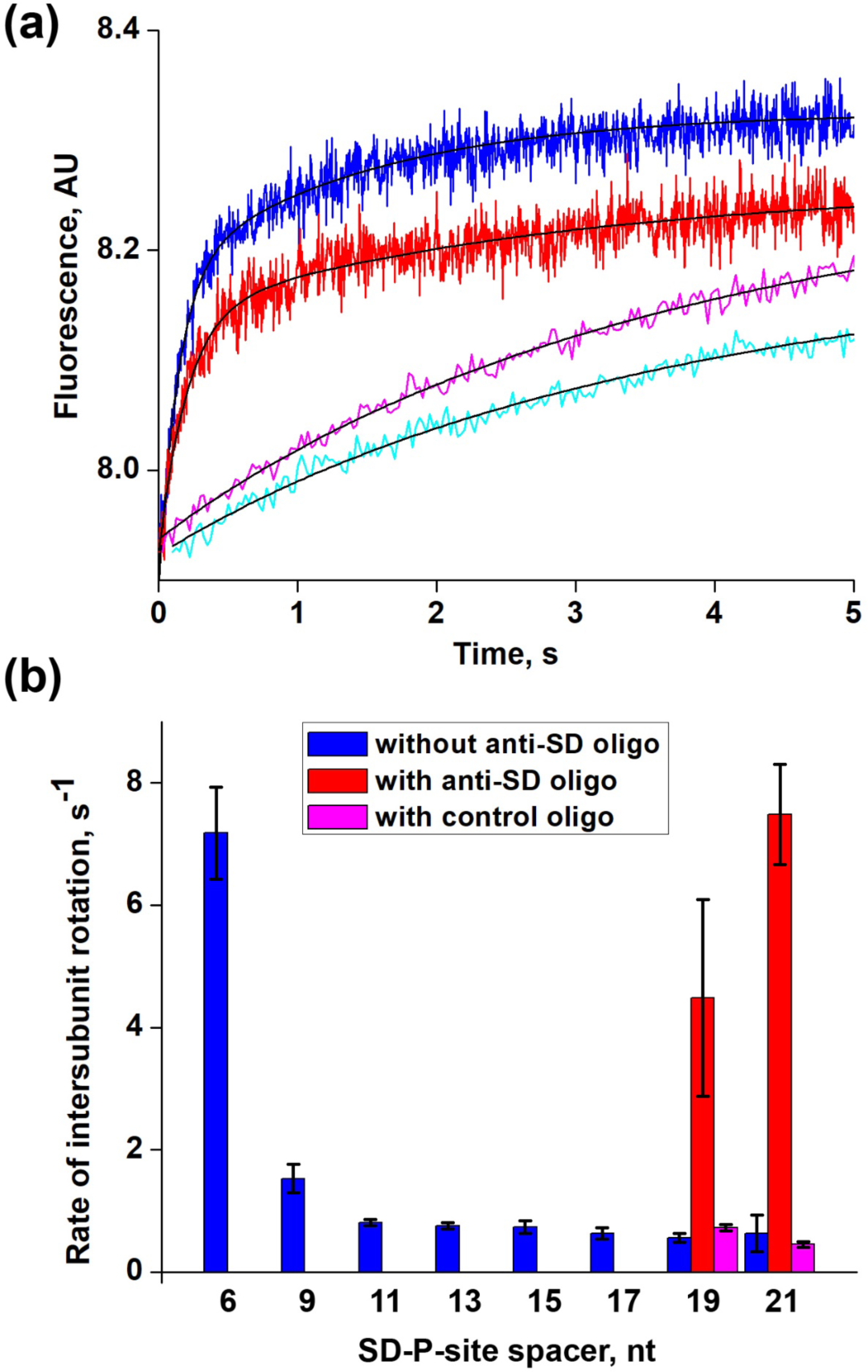
Kinetics of intersubunit rotation coupled to translocation of mRNAs with different spacing between the SD and P-site codon (UAC). Pre-translocation S6-Alexa488 (donor)/L9-Alexa568 (acceptor) ribosomes containing deacylated tRNA^Tyr^ in the P site and N-acetyl-Phe-tRNA^Phe^ in the A site were rapidly mixed with EF-G and GTP. Pre-translocation ribosomes were programmed by mRNAs containing a AAGGA SD sequence. Spacing between the SD and P-site (UAC) codon varied from 6 to 21 nucleotides as indicated. **(a)** Kinetics of intersubunit rotation followed by FRET (acceptor fluorescence) in ribosomes programmed by mRNA with 6-nucleotide long SD-UAC spacer (blue); mRNA with 21-nucleotide long SD-UAC spacer (cyan); mRNA with 21-nucleotide long SD-UAC spacer annealed to either an anti-SD DNA oligo (red) or a control DNA oligo (magenta). Double-exponential fits are black lines. **(b)** Bar graphs showing rates of intersubunit rotation (*k*_*1*_ of double-exponential fit) coupled to translocation of mRNAs with various SD-UAC codon spacing (blue). Rates measured in ribosomes programmed with mRNAs annealed to either an anti-SD DNA oligo or a control DNA oligo are shown in red and magenta, respectively.

Ribosomes programmed by mRNA with an 6 nucleotide-long spacer between the AAGGA SD sequence and P-site (UAC) codon translocated rapidly (*k*_*1*_=7.2 ± 0.8 s^-1^). Extending spacing between the AAGGA SD sequence and UAC codon to 9 nucleotides substantially slowed the rate of reverse intersubunit rotation coupled to mRNA translocation to 1.5 s^-1^ (Fig. 3b). Even lower rates of 0.6-0.8 s^-1^ were observed for mRNAs with a 11, 13, 15, 17, 19 or 21-nucleotide long spacer between the AAGGA SD sequence and P-site (UAC) codon (Fig 3b, Suppl. Table 1). Extending the SD-P-site spacer also decreased the amplitude of fluorescence change (Fig. 3a) likely indicating lower stability of pre-translocation complexes programmed by these mRNAs (Fig. 1 and Suppl. Fig. 1). These results suggest that extending the spacing between the SD sequence and P-site codon inhibits the intersubunit rotation coupled to mRNA translocation.

A possible alternative interpretation of the kinetics experiments is that extending the spacing between the AAGGA SD sequence and UAC codon beyond 9 nucleotides disrupts SD-aSD interactions. To measure the rate of translocation in the absence of SD-aSD interactions, DNA oligo complementary to SD sequence and 15 nucleotides upstream of SD was added to pre-translocation ribosomes programmed with an mRNA, which contained either 19 or 21 nucleotide-long SD-P-site spacer. Anti-SD DNA oligo is expected to compete with aSD sequence of 16S rRNA and thus sequester SD sequence into DNA-RNA duplex. Annealing anti-SD DNA oligo dramatically increased the rate of ribosome translocation from 0.6 to 4.5 s^-1^ for 19-nucleotide spacing, and from 0.6 to 7.5 s^-1^ for 21-nucleotide spacing between SD and UAC codon (Fig. 3, Suppl. Table 1).

As a negative control, instead of anti-SD DNA oligo, 20-nuleotide long DNA oligo complementary to mRNA sequence upstream of SD (but not SD sequence itself) was added to pre-translocation complexes with 19 or 21 spacers between SD and UAC codon. In contrast to anti-SD DNA oligo, annealing control DNA oligos did not increase the rate of translocation (Fig. 3 and Suppl. Table 1). Our results suggest that SD-aSD interactions remain intact and slow down translocation even when the spacer between SD and UAC codon is 19 or 21-nucleotides long. Remarkably, ribosomes programmed by mRNA with 6-nucleotide long spacer between SD and UAC codon and ribosomes programmed by mRNA with 21-nucleotide long spacer, in which SD-aSD interactions were disrupted by the anti-SD DNA oligo, translocated essentially at the same rate. Hence, when spacing between SD and P-site codon is short, mRNA translocation is equally fast as translocation in the absence of SD-aSD interactions. However, extending the SD-P site spacer slows down translocation until SD-aSD helix is unwound.

In the context of a 6-nucleotide spacer between SD and UAC (P-site) codon, replacing the AAGGA SD sequence with the stronger SD sequence AAGGAGGU, decreased the rate of translocation from 7.2 (AAGGA SD) to 1.5 s^-1^ (AAGGAGGU SD) (Fig. 4, Suppl. Table 2). In the presence of the stronger SD sequence AAGGAGGU, extending SD-P-site spacer from 6 to 9 or 11 nucleotides decreased the rate of translocation from 1.5 to 0.6 or 0.8 s^-1^, respectively (Fig. 4, Suppl. Table 2). Hence, increasing the strength of SD-aSD interactions slows down ribosome translocation.

**Figure 4.**
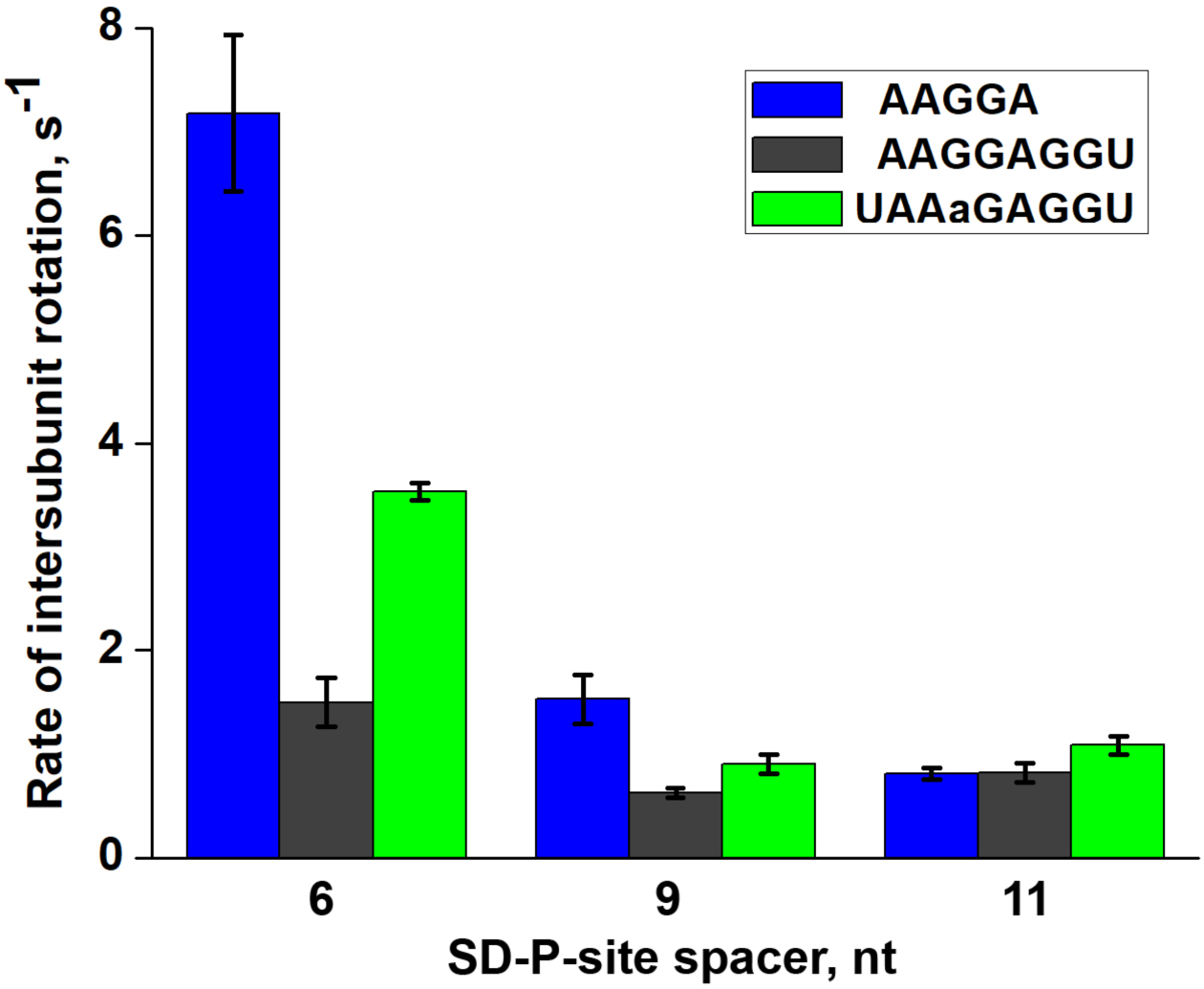
Kinetics of intersubunit rotation coupled to translocation of mRNAs containing different SD sequences. Pre-translocation S6-Alexa488 (donor)/L9-Alexa568 (acceptor) ribosomes containing deacylated tRNA^Tyr^ in the P site and N-acetyl-Phe-tRNA^Phe^ in the A site were rapidly mixed with EF-G and GTP. Pre-translocation ribosomes were programmed by mRNAs containing either AAGGA (blue), AAGGAGGU (black) or UAAaGAGGU (green) SD sequence. Spacing between the SD and P-site (UAC) codon varied from 6 to 11 nucleotides as indicated.

Consistent with this conclusion, introduction of a mismatch into SD-aSD duplex in the context of 6-nucleotide spacer between SD and UAC (P-site) codon, increased the rate of translocation from 1.5 (with AA<u>G</u>GAGGU SD) to 3.5 s^-1^ (with UAA<u>a</u>GAGGU SD) (Fig. 4, Suppl. Table 3). In the presence of UAA<u>a</u>GAGGU SD sequence, extending SD-P-site spacer from 6 to 9 or 11 nucleotides decreased the rate of translocation from 3.5 to 0.9 or 1.1 s^-1^, respectively (Fig. 4, Suppl. Table 3).

Our kinetic data indicate that regardless of SD sequence, extending the spacing between the SD and P-site codon slows down translocation. However, in the presence of the stronger SD sequence, variation in SD-P-site spacer length has a smaller effect on translocation rate. In contrast to AAGGA SD sequence, the stronger SD sequence AAGGAGGU substantially inhibits translocation even when the spacer between SD and P-site codon is as short as 6 nucleotides.

### Extending the spacing between the SD and P-site codon inhibits mRNA translocation

A possible caveat in aforementioned kinetic experiments is that we measured the rate of mRNA translocation indirectly by measuring the rate of reverse intersubunit rotation of the ribosome from the rotated into non-rotated conformation, which has been previously shown to be coupled to mRNA translocation [26]. To determine the rate of mRNA translocation directly, we measured fluorescence quenching of a fluorescein dye attached to the 3’ end of short model mRNAs as they move within the ribosome. This fluorescence quenching translocation assay was extensively used before [26, 32-34].

We tested the effect of the length of the spacer between the SD sequence and P-site (AUG) codon on the rate of translocation in the presence of a strong (AAGGAGGU) SD sequence. We also measured the rate of translocation of a leaderless mRNA (5’ AUG UAC AAA GUA UAA 3’) that does not contain a SD sequence. When pre-translocation ribosomes, which were assembled with fluorescein-labeled mRNA, deacylated tRNA^Met^ in the P site and N-acetyl-Tyr-tRNA^Tyr^ in the A site, were mixed with EF-G•GTP using a stopped-flow apparatus, rapid quenching of fluorescein fluorescence was observed, indicative of mRNA translocation (Fig. 5). Consistent with published reports [26, 29-31] and our S6Alexa488/L9Alexa586 kinetic FRET experiments (Fig. 3), kinetic traces of mRNA translocation were best fitted to a double-exponential function.

**Figure 5.**
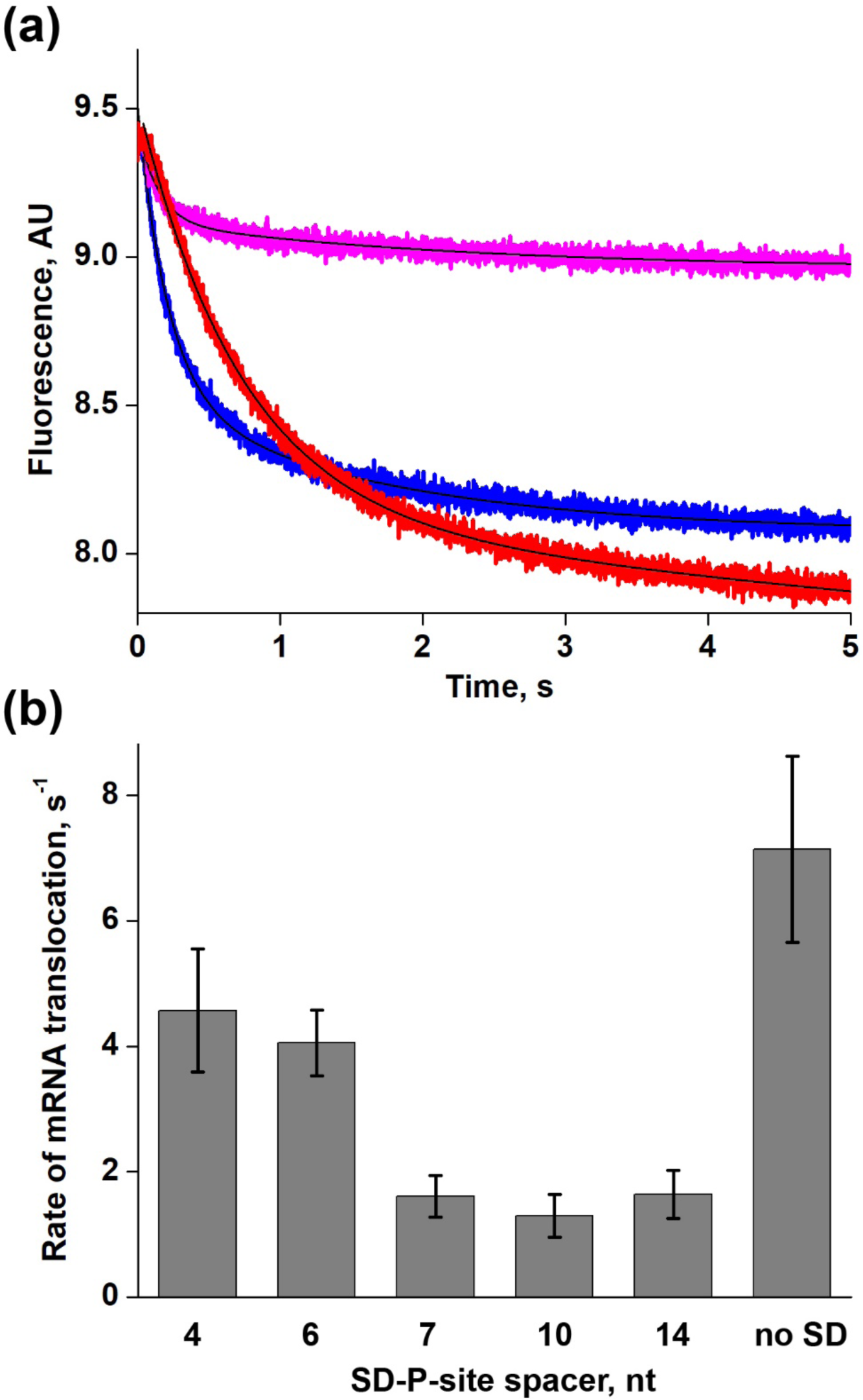
Kinetics of mRNA translocation measured by quenching of fluorescein attached to the 3’ end of mRNA. Pre-translation ribosomes containing deacylated tRNA^Tyr^ in the P site and N-acetyl-Phe-tRNA^Phe^ in the A site were rapidly mixed with EF-G and GTP. Pre-translocation ribosomes were programmed by fluorescein-labeled mRNAs. All mRNAs contained a AAGGAGGU SD sequence except for the “no SD” leaderless mRNA, which lacked the SD. Spacing between the SD and P-site (UAC) codon varied from 4 to 14 nucleotides as indicated. **(a)** Kinetics of fluorescein quenching corresponding to translocation of mRNA with 4 nucleotide-long spacing between the SD and UAC codon (blue), mRNA with 10 nucleotide-long spacing between the SD and UAC codon (red) and leaderless mRNA lacking the SD (magenta). Double-exponential fits are black lines. **(b)** Rates of mRNA translocation (*k*_*1*_ of double-exponential fit).

Similarly fast translocation was observed for the mRNAs with the 4 and 6 nucleotide spacer between the SD and P-site codon AUG as faster rate constants *k*_*1*_ of the bi-phasic fit were 4.6 and 4.1 s^-1^, respectively (Fig. 5 and Suppl. Table 4). By contrast, 2.5-3 fold decreases in translocation rates were observed for mRNAs having a 7, 10 or 14-nucleotide spacer (Fig. 5, Suppl. Table 4).

Amplitude of fluorescein quenching observed for leaderless mRNA that lacks the SD sequence was significantly lower than the amplitude observed for mRNAs containing the SD (Fig. 5). This decrease in amplitude is likely due to markedly lower stability of mRNA-ribosome complex in the absence of the SD. Nevertheless, leaderless mRNA translocated slightly faster (with the rate constant *k*_*1*_ of 7.1 s^-1^) than mRNAs with 4 and 6-nucleotide long spacers between the SD and P-site codon (Fig. 5 and Suppl. Table 4). Taken together, our kinetics measurements show that when the spacer between the SD and P site codon extends beyond 4-6 nucleotides, the SD-aSD helix appreciably slows down translocation.

## Discussion

Our experiments revealed that the effect of SD sequence on ribosome translocation is variable and depends on spacing between the SD and P-site codon. In most of our experiments (except the mRNA derived from m301 mRNA, which contained strong SD sequence AAGGAGGU), when the SD-P-site spacer was 4-6 nucleotides long, the rate of translocation was similar to the rate observed in the absence of SD-aSD interactions (Fig. 3 and 5). This result is consistent with experiments performed by Borg and Ehrenberg showing that three different SD-like sequences positioned six nucleotides upstream of P-site codon had no effect on translocation rate [8].

Our data also indicate that extending spacing between the SD sequence and P-site codon beyond 6 nucleotides substantially inhibits ribosome translocation (Fig. 3-5). Such SD-aSD interactions can result in 5-10 fold reduction of ribosome translocation rate when compared to the rate observed in the absence of SD-aSD interactions. These observations are consistent with previous *in vitro* single-molecule experiments demonstrating that SD sequences slows down translocation by 3-4 fold when the spacer between SD sequence and P-site codon is 9-15 nucleotides long [6].

Only at 6 nucleotide-long SD-P-site spacing, the inhibitory effect of SD-aSD interactions inversely correlated with the rate of translocation: strongest SD sequence AAGGAGGU slowed down translocation to the largest degree (Fig. 4). By contrast, mRNAs that contained AAGGA, AAGGAGGU or UAAaGAGGU SD sequences 7-14 nucleotides upstream of the P site were translocated at similar rates (Fig. 3-5) suggesting that at least under experimental conditions used in our *in vitro* studies, the effect of SD sequence on translocation rate primarily depends on the SD-P-site spacing and not the strength of SD-aSD interactions. In model mRNAs used in this study, the SD-P-site spacer was extended by inserting intrinsically unstructured CA repeats. Unexplored in our study, nucleotide composition and basepairing potential of the SD-P-site spacer sequence may also affect the rate of translocation thus increasing variability of the effect of SD sequence on ribosome translocation.

Our kinetic experiments performed in the presence of anti-SD DNA oligo annealed to mRNA containing a short SD sequence AAGGA (Fig. 3) indicate that at least under conditions used in our *in vitro* experiments, SD-aSD interactions can be retained when the spacer between SD and P site codon is ≤ 21-nucleotide long. Hence, upon progressive movement of the ribosome away from the SD sequence, mRNA likely undergoes inchworm rearrangement inside the ribosome. This idea is supported by *in vivo* ribosome profiling data demonstrating that SD-like sequences produce ribosome-protected footprints that are up to 15 nucleotide longer than typical 28-29 nt-long ribosome protected mRNA fragments [4]. Furthermore, these longer ribosome-protected fragments are asymmetrically elongated at the 3’ end, consistent with inchworm-like rearrangement of mRNA anchored by SD sequence to the 30S subunit.

Published structural studies provide possible clues for the mechanism of SD-induced inhibition of translocation. X-ray crystal structures of several ribosome-mRNA complexes with different SD-P-site spacing indicated that during the first of several elongation cycles, SD-aSD helix undergoes rotational movement and shifts away from the intersubunit interface toward the solvent side of the 30S subunit closer to the 30S head domain [35, 36]. In the crystal structure of the ribosome with 9 nucleotide spacer between the SD and P-site codon, three nucleotides of the spacer adjacent to SD were stacked on top of basepairs formed by SD and 16S rRNA thus extending SD-aSD helix [36]. Extension of the spacer between SD and P-site codon may also lead to other structural changes that are yet to be visualized by X-ray crystallography and cryo-EM. We hypothesize that mRNA rearrangements resulting from the extension of SD-P-site spacer cause inhibition of mRNA translocation. One possibility is that these structural rearrangements interfere with the rotational movement of the 30S head domain that is thought to accompany mRNA translocation [37].

Our work provides new insights into the roles of SD sequences in translation initiation and elongation. The inhibition of translocation by SD sequences detected in our experiments may affect the transition from initiation to elongation phase of protein synthesis [38] and at least partially be responsible for the slower translation of the first several codons in ORFs that has been observed in several previous studies [8, 39].

Our data also support the idea that intragenic SD sequences can regulate translation elongation. Although published ribosome profiling studies produced conflicting data regarding the ability of SD-like sequences to induce translational pauses *in vivo* [3, 7], SD-aSD interactions likely slow down ribosome translocation to a certain degree in live cells. It is possible the 5-10 fold reduction of ribosome translocation rate observed in our experiments is not sufficient to induce lengthy translation pauses that can be detected by ribosome profiling approach. It is also possible that the effect of SD-aSD interactions on ribosome translocation is exacerbated under *in vitro* conditions because concentration of free Mg2+ ions, which likely stabilize SD-aSD interactions, is lower in live cells. Hence, in comparison to *in vitro* experiments, SD-induced inhibition of translocation may less prominent in live cells. Nevertheless, conservation of intragenic SD-like sequences between divergent bacterial species [2] support the idea that SD-mediated regulation of translation elongation is functionally important *in vivo*.

Toeprinting (Fig. 1-2) and kinetic (Fig. 3-5) data also provide insights into the mechanism of SD-stimulated programmed ribosome frameshifting (PRF) in bacteria. Our results suggest that extending SD-P-site spacer beyond 9 nucleotides destabilizes mRNA-ribosome interactions and promotes mRNA back-slippage. Consistent with these observations, the optimal spacing between the P site codon and SD sequences for stimulation of -1 PRF in *E.coli dnaX* mRNA was shown to be 10-14 nucleotides [10]. Therefore, SD sequences likely stimulate -1 PRF by (i) slowing down ribosome translocation and (ii) destabilizing of codon-anti-codon interactions in the “0” reading frame.

## Materials and Methods

### Materials and sample preparation

tRNA^Phe^ and tRNA^Tyr^ were purchased from Chemblock. MRE600 *E.coli* 70S ribosomes, 6-histidine-tagged EF-G and aminoacylated tRNAs were prepared as previously described [18, 24]. Using the QuickChange Site-Directed Mutagenesis System (Agilent, Santa Clare, CA), DNA templates encoding mRNA variants for toeprinting and kinetic experiments were derived from pFK301 plasmid, which encodes m301 mRNA (5’ GUAAAGUGUCAUAGCACCAACUGUUAAUUAAAUUAAAUUAAA<u>AAGGA</u>AAUAAUGUU<u>UAC</u>U UUGUAAAAUCUACUGCUGAACUCGCUGCACAAAUGGCUAAACUGAAUGGCAAUAAAGGUUUUUCUUCUGAAGAUAAAG 3’; the SD sequence and a UAC codon are underlined [19]). Sequences of all mRNA variants are shown in Suppl. Table 4.

### Fluorescently-labeled mRNAs and ribosomes

S6-Alexa488/L9-Alexa568 ribosomes were prepared as previously described [24, 27]. Single cysteine mutants of the small ribosomal subunit protein S6 and large ribosomal subunit protein L9 were labeled using maleimide derivatives of Alexa488 and Alexa568, respectively (Thermo Fisher Scientific, Waltham, MA). Labeled S6 and L9 proteins were incorporated into ΔS6 30S and ΔL9 50S subunits, respectively, by partial reconstitution as previously described [24, 27].

Fluorescein labeled mRNAs were synthesized by Integrated DNA Technologies (Coralville, IA). mRNA variants were derived from the mRNA with 4-nucleotide spacer between SD and AUG codon [33] (5’ GGC<u>AAGGAGGU</u>AAAA<u>AUG</u>UACAAAGUAUAA 3’ Fluorescein; SD sequence and AUG codon are underlined): 5’ GGC<u>AAGGAGGU</u>ACACAA<u>AUG</u>UACAAA 3’Fluorescein (6 nt spacer); 5’ GGC<u>AAGGAGGU</u>ACACAAA<u>AUG</u>UACAAA 3’Fluorescein (7 nt spacer); 5’ GGC<u>AAGGAGGU</u>ACAACACAAA<u>AUG</u>UACAAA 3’Fluorescein (10 nt spacer); 5’ GGC<u>AAGGAGGU</u>AACAACACAAACAA<u>AUG</u>UACAAA 3’Fluorescein (14 nt spacer). We also measured the rate of translocation of leaderless mRNA (5’ <u>AUG</u>UACAAAGUAUAA 3’ Fluorescein) that does not contain SD sequence.

### Toeprinting experiments

Toeprinting experiments were performed in polyamine buffer (30 mM HEPES•KOH, pH 7.5, 70 mM NH_4_Cl, 6 mM MgCl_2_, 2 mM spermidine, 0.1 mM spermine) as previously described [18, 21]. Complexes in Fig. 1 were assembled by incubating 70S ribosomes (1 μM) with deacylated tRNA^Tyr^ (2 μM) and mRNA (2 μM) preannealed to [^32^P]-labeled primer [18] for 15 minutes at 37 °C. The pre-translocation complexes in Suppl. Fig. S1 were assembled by incubating 70S ribosomes (1 μM) with deacylated tRNA^Tyr^ (2 μM) and mRNA (2 μM) preannealed to [^32^P]-labeled primer [18] for 15 minutes at 37 °C. Then, 1.5 µM *N*-acetyl-Phe-tRNA^Phe^ was added to P-site tRNA-bound complexes followed by incubation at 37 °C for 20 minutes. The pre-translocation complexes in Fig. 2 were assembled similarly except that the A site was filled with 2 µM deacylated tRNA^Phe^ added to P-site tRNA-bound complexes. Translocation was carried out by the incubation of pre-translocation complexes (0.5 μM) with 1.5 μM EF-G in the presence of 0.5 mM GTP for 10 minutes at 37 °C.

### Stopped-flow measurements of pre-steady-state translocation kinetics

Kinetics of ribosome intersubunit rotation coupled to mRNA translocation were measured as previously described [26]. Pre-translocation complexes were constructed by the incubation of S6-Alexa488/L9-Alexa568 70S ribosomes (1.4 µM) with mRNA 2.8 µM and deacylated tRNA^Tyr^ (4 µM) in polyamine buffer (30 mM HEPES•KOH, pH 7.5, 150 mM NH_4_Cl, 6 mM MgCl_2_, 2 mM spermidine, 0.1 mM spermine, 6 mM β-mercaptoethanol) for 15 minutes at 37 °C, followed by an incubation with *N*-acetyl-Phe-tRNA^Phe^ (2.8 µM) for 30 minutes at 37 °C. In experiments shown in Fig. 3, 28 µM anti-SD (5’-TCCTTTTTAATTTAATTTAA-3’) or control DNA (5’-ATTTAATTTAATTAACAGTT-3’) oligos were annealed to mRNA after complex assembly for 15 min at 37 °C. Anti-SD and control oligo were designed to have similar affinity to mRNA. Pre-translocation ribosomes were mixed with EF-G and GTP (or GTP analogues) using an Applied Photophysics stopped-flow fluorometer. Final concentrations after mixing were: 35 nM ribosomes, µM EF-G, 0.5 mM GTP and 14 µM DNA oligo (if oligo was added to pre-translocation ribosomes). Alexa488 was excited at 490 nm and changes in FRET (Alexa568 emission) were detected using a 590 nm long-pass filter. All stopped-flow experiments were done at 23°C; monochromator slits were adjusted to 9.3 nm. Increases in FRET (Alexa568 fluorescence) in kinetics traces were best fitted to the sum of two exponentials (y=y_0_+A_1_*exp(-k_1_*t)+A_2_*exp(-k_2_*t)), corresponding to the apparent rate constants *k*_*1*_ and *k*_*2*_.

Kinetics of mRNA translocation were measured by fluorescein quenching as previously described with minor modifications [26, 32, 34]. Pre-translocation complexes were constructed by the incubation of 70S ribosomes (1 µM) with fluorescein-labeled mRNA (0.85 µM) and deacylated tRNA^Met^ (2 µM) in polyamine buffer (30 mM HEPES•KOH, pH 7.5, 150 mM NH_4_Cl, 6 mM MgCl_2_, mM spermidine, 0.1 mM spermine) for 15 minutes at 37 °C, followed by an incubation with *N*-acetyl-Tyr-tRNA^Tyr^ (1.5 µM) for 30 minutes at 37 °C. Pre-translocation ribosomes were mixed with EF-G and GTP using an Applied Photophysics stopped-flow fluorometer (Leatherhead, Surrey, UK). Final concentrations after mixing were: 35 nM ribosomes, 1 µM EF-G and 0.5 mM GTP. Fluorescein was excited at 490 nm and fluorescence emission was detected using a 515 nm long-pass filter. All stopped-flow experiments were done at 23°C; monochromator slits were adjusted to 9.3 nm. Translocation of the mRNA resulted in a partial quenching of fluorescein coupled to the 3’ end of the mRNA (8). Time traces were analyzed using Origin (OriginLab Corporation, Northampton, MA). As reported previously [26, 30, 31, 33], the kinetics of mRNA translocation are clearly biphasic and are best fitted to the sum of two exponentials (y=y_0_+A_1_*exp(-k_1_*t)+A_2_*exp(-k_2_*t)), corresponding to the apparent rate constants *k*_*1*_ and *k*_*2*_.

## Abbreviations used

SD: Shine-Dalgarno;
aSD: anti-Shine-Dalgarno;
ORFs: open reading frames;
PRF: programmed ribosome frameshifting;
RF2: release factor 2;
FRET: Förster resonance energy transfer;
EF-G: elongation factor G.

## Acknowledgements

These studies were supported by grant from the US National Institute of Health R01GM099719 (to D.N.E.). We thank Andrei Korostelev for his comments on the manuscript.

## Supplementary Materials

**Supplementary Figure 1.**
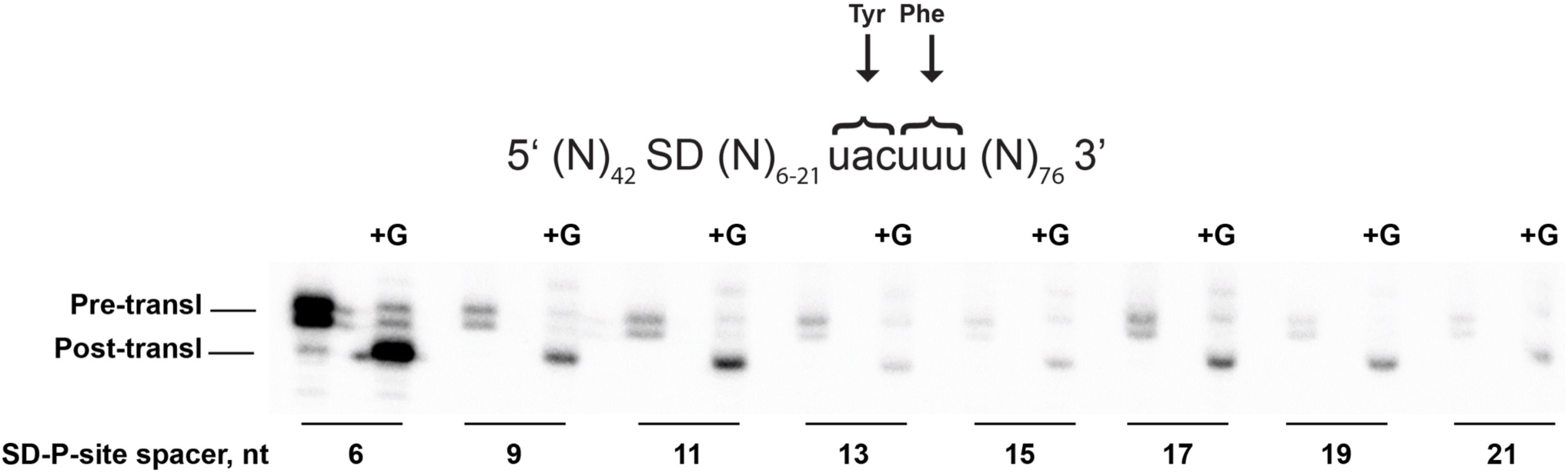
Extending spacing between the SD sequence and the P site codon destabilizes mRNA-ribosome interactions. A pre-translocation complex (“Pre-transl”) was made by binding deacylated tRNA^Tyr^ to the P site and N-acetyl-Phe-tRNA^Phe^ to the A site in the presence of an mRNA. The spacer between the AAGGA SD sequence and UAC (P-site) codon varied between 6 and 21 nucleotides in length as indicated. The position of the ribosome along the mRNA was mapped by toeprinting. Pre-translocation complexes were incubated with EF-G and GTP to induce mRNA translocation (+G lanes). “Post-transl” toeprint bands correspond to the product of translocation.

**Supplementary Table 1.** Rates of intersubunit rotation coupled to translocation of mRNAs containing an AAGGA SD sequence. Pre-translocation S6-Alexa488 (donor)/L9-Alexa568 (acceptor) ribosomes containing deacylated tRNA^Tyr^ in the P site and N-Ac-Phe-tRNA^Phe^ in the A site were rapidly mixed with EF-G and GTP (Fig. 3). Pre-translocation ribosomes were programmed by mRNAs containing a AAGGA SD sequence. Spacing between the SD and P-site (UAC) codon varied from 6 to 21 nucleotides as indicated. About ten traces were acquired for each experiment. *k*_*1*_ and *k*_*2*_ are rates constants of double-exponential fits. Rate constants averaged from two to four independent experiments are shown.

| Spacer between the SD and UAC codon, nt | $k_1$ (s<sup>-1</sup>) | $k_2$ (s<sup>-1</sup>) |
| --- | --- | --- |
| 6 | $7.2 \pm 0.8$ | $0.9 \pm 0.22$ |
| 9 | $1.5 \pm 0.2$ | $0.4 \pm 0.11$ |
| 11 | $0.8 \pm 0.1$ | $0.2 \pm 0.01$ |
| 13 | $0.8 \pm 0.1$ | $0.2 \pm 0.02$ |
| 15 | $0.7 \pm 0.1$ | $0.2 \pm 0.05$ |
| 17 | $0.6 \pm 0.1$ | $0.2 \pm 0.02$ |
| 19 | $0.6 \pm 0.1$ | $0.1 \pm 0.03$ |
| 19 (with anti-SD oligo) | $4.5 \pm 1.6$ | $0.6 \pm 0.24$ |
| 19 (with control oligo) | $0.7 \pm 0.1$ | $0.1 \pm 0.01$ |
| 21 | $0.6 \pm 0.3$ | $0.1 \pm 0.05$ |
| 21 (with anti-SD oligo) | $7.5 \pm 0.8$ | $1.3 \pm 0.41$ |
| 21(with control oligo) | $0.5 \pm 0.1$ | $0.1 \pm 0.01$ |

**Supplementary Table 2.**
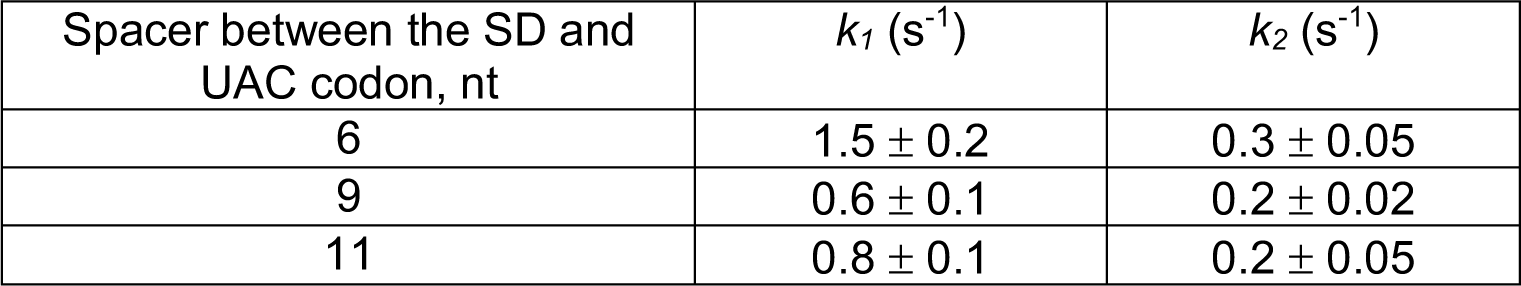
Rates of intersubunit rotation coupled to translocation of mRNAs with AAGGAGGU SD sequence. Pre-translocation S6-Alexa488 (donor)/L9-Alexa568 (acceptor) ribosomes containing deacylated tRNA^Tyr^ in the P site and N-Ac-Phe-tRNA^Phe^ in the A site were rapidly mixed with EF-G and GTP (Fig. 4). Pre-translocation ribosomes were programmed by mRNAs containing a AAGGAGGU SD sequence. Spacing between the SD and P-site (UAC) codon varied from 6 to 21 nucleotides as indicated. About ten traces were acquired for each experiment. *k*_*1*_ and *k*_*2*_ are rates constants of double-exponential fits. Rate constants averaged from two to four independent experiments are shown.

**Supplementary Table 3.** Rates of intersubunit rotation coupled to translocation of mRNAs with UAAaGAGGU SD sequence. Pre-translocation S6-Alexa488 (donor)/L9-Alexa568 (acceptor) ribosomes containing deacylated tRNA^Tyr^ in the P site and N-Ac-Phe-tRNA^Phe^ in the A site were rapidly mixed with EF-G and GTP (Fig. 4). Pre-translocation ribosomes were programmed by mRNAs containing a UAAaGAGGU SD sequence. Spacing between the SD and P-site (UAC) codon varied from 6 to 21 nucleotides as indicated. About ten traces were acquired for each experiment. *k*_*1*_ and *k*_*2*_ are rates constants of double-exponential fits. Rate constants averaged from two to four independent experiments are shown.

| Spacer between the SD and UAC codon, nt | $k_1$ (s<sup>-1</sup>) | $k_2$ (s<sup>-1</sup>) |
| --- | --- | --- |
| 6 | $3.6 \pm 0.1$ | $0.5 \pm 0.03$ |
| 9 | $0.9 \pm 0.1$ | $0.2 \pm 0.03$ |
| 11 | $1.1 \pm 0.1$ | $0.3 \pm 0.02$ |

**Supplementary Table 4.**
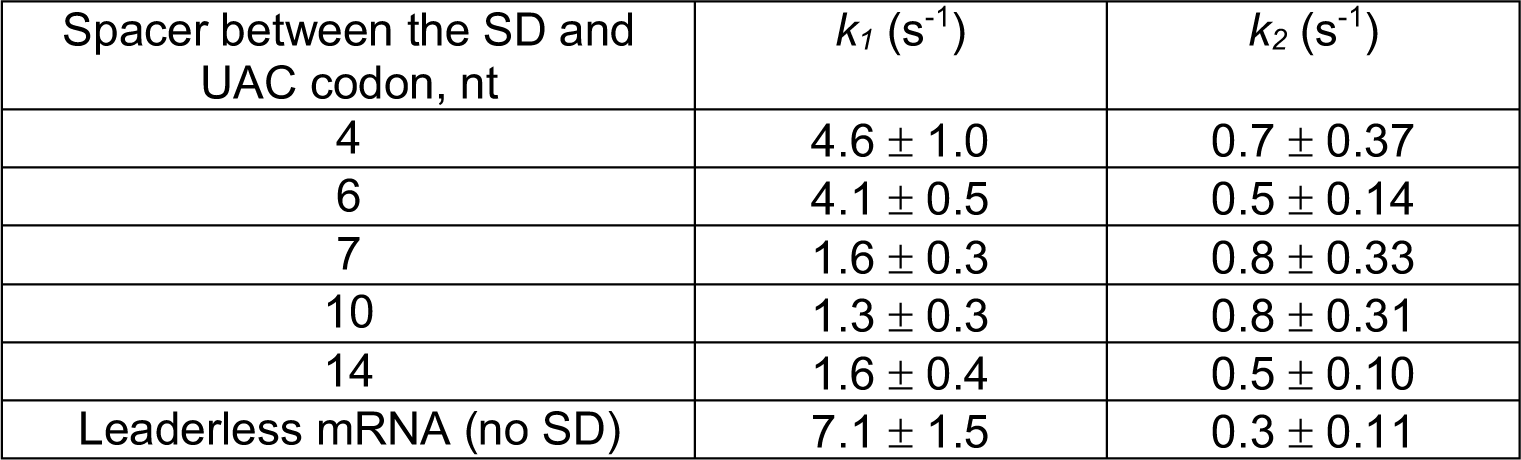
Rates of mRNA translocation determined by quenching of fluorescein attached to the 3’ end of mRNA. Pre-translation ribosomes containing deacylated tRNA^Tyr^ in the P site and N-Ac-Phe-tRNA^Phe^ in the A site were rapidly mixed with EF-G and GTP. Pre-translocation ribosomes were programmed by fluorescein-labeled mRNAs. All mRNAs contained a AAGGAGGU SD sequence except for the “no SD” leaderless mRNA, which lacked the SD. Spacing between the SD and P-site (UAC) codon varied from 4 to 14 nucleotides as indicated. About ten traces were acquired for each experiment. *k*_*1*_ and *k*_*2*_ are rates constants of double-exponential fits. Rate constants averaged from two to four independent experiments are shown.

**Supplementary Table 5.**
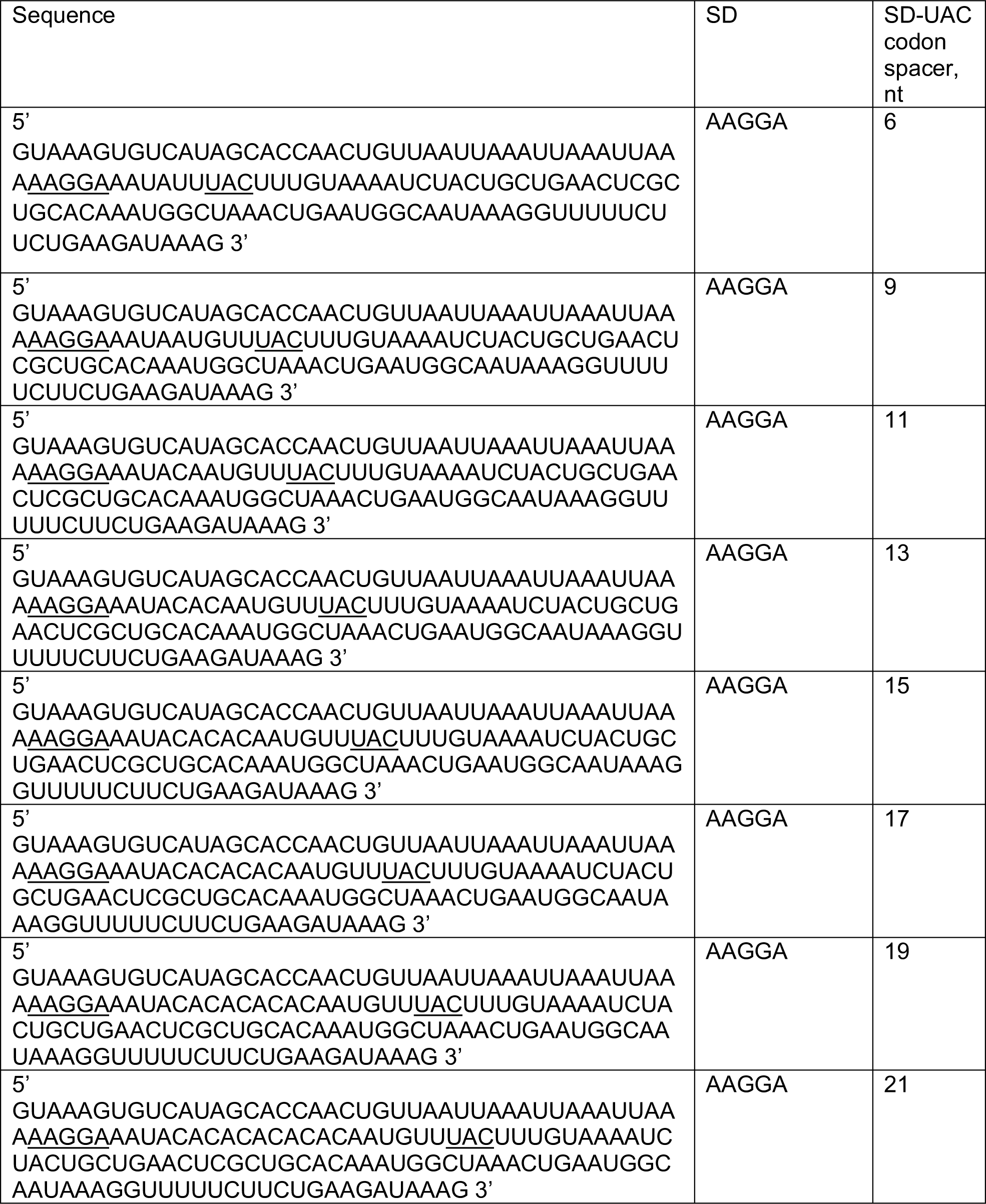
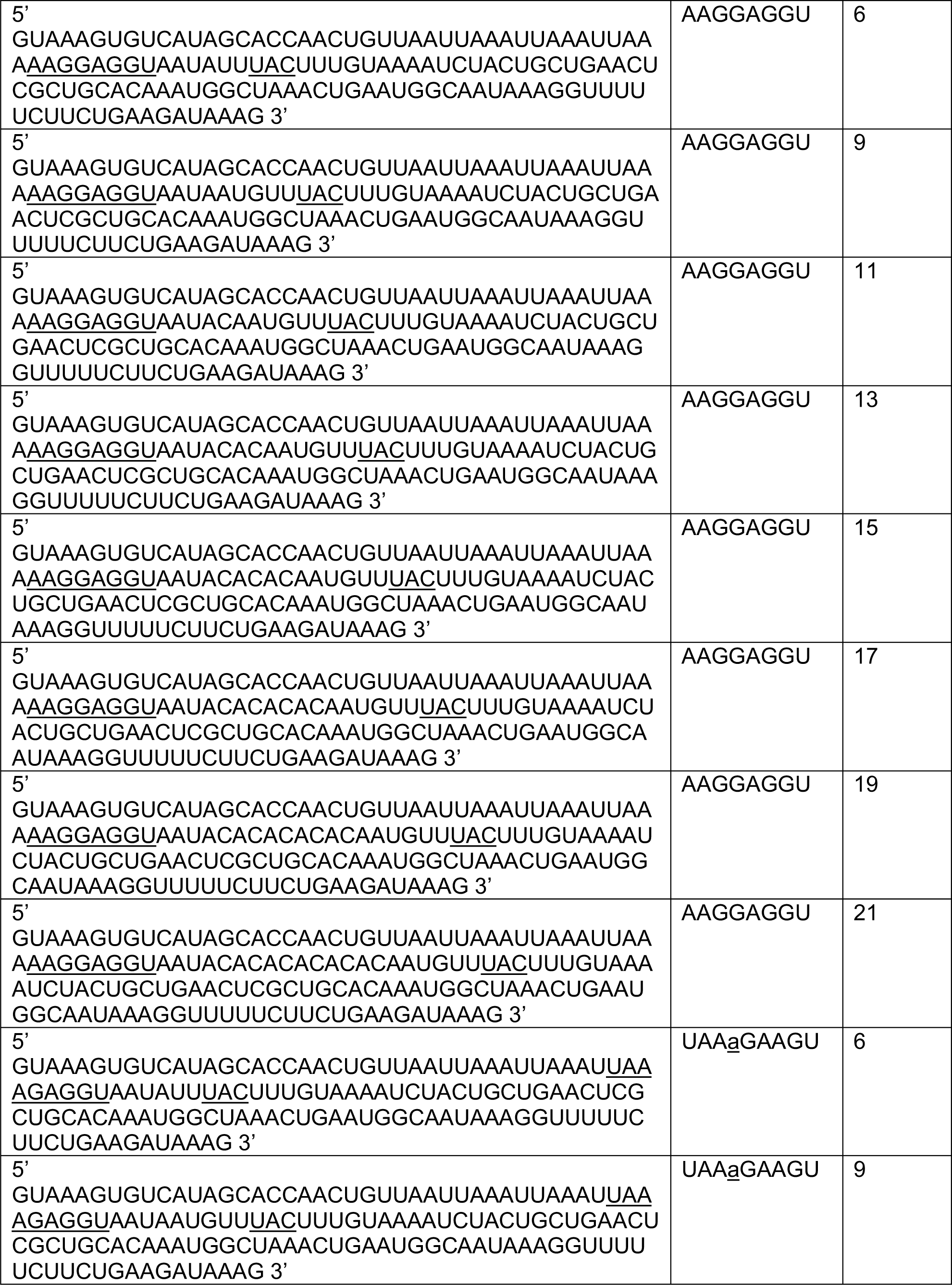
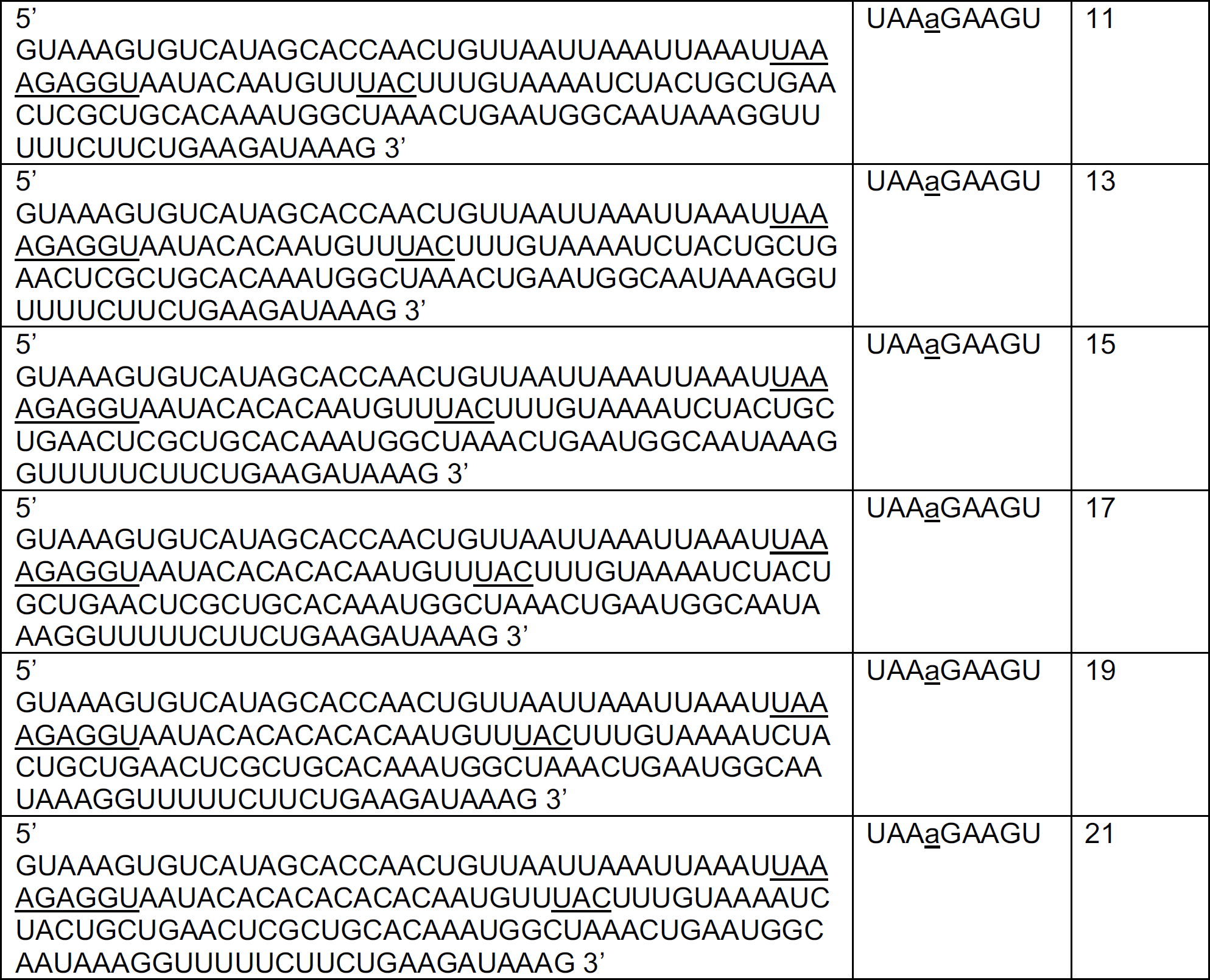
Sequences of mRNAs used in toeprinting and S6-Alexa488/L9-Alexa568 kinetic experiments.

## References

[1] Steitz JA, Jakes K. How ribosomes select initiator regions in mRNA: base pair formation between the 3’ terminus of 16S rRNA and the mRNA during initiation of protein synthesis in Escherichia coli. Proc Natl Acad Sci U S A. 1975;72:4734–8.

[2] Saito K, Green R, Buskirk AR. Translational initiation in E. coli occurs at the correct sites genome-wide in the absence of mRNA-rRNA base-pairing. Elife. 2020;9.

[3] Li GW, Oh E, Weissman JS. The anti-Shine-Dalgarno sequence drives translational pausing and codon choice in bacteria. Nature. 2012;484:538–41.

[4] O’Connor PB, Li GW, Weissman JS, Atkins JF, Baranov PV. rRNA:mRNA pairing alters the length and the symmetry of mRNA-protected fragments in ribosome profiling experiments. Bioinformatics. 2013;29:1488–91.

[5] Wen JD, Lancaster L, Hodges C, Zeri AC, Yoshimura SH, Noller HF, et al. Following translation by single ribosomes one codon at a time. Nature. 2008;452:598–603.

[6] Chen J, Petrov A, Johansson M, Tsai A, O’Leary SE, Puglisi JD. Dynamic pathways of -1 translational frameshifting. Nature. 2014;512:328–32.

[7] Mohammad F, Woolstenhulme CJ, Green R, Buskirk AR. Clarifying the Translational Pausing Landscape in Bacteria by Ribosome Profiling. Cell Rep. 2016;14:686–94.

[8] Borg A, Ehrenberg M. Determinants of the rate of mRNA translocation in bacterial protein synthesis. J Mol Biol. 2015;427:1835–47.

[9] Spanjaard RA, van Duin J. Translation of the sequence AGG-AGG yields 50% ribosomal frameshift. Proc Natl Acad Sci U S A. 1988;85:7967–71.

[10] Larsen B, Wills NM, Gesteland RF, Atkins JF. rRNA-mRNA base pairing stimulates a programmed -1 ribosomal frameshift. J Bacteriol. 1994;176:6842–51.

[11] Atkins JF, Baranov PV, Fayet O, Herr AJ, Howard MT, Ivanov IP, et al. Overriding standard decoding: implications of recoding for ribosome function and enrichment of gene expression. Cold Spring Harb Symp Quant Biol. 2001;66:217–32.

[12] Caliskan N, Peske F, Rodnina MV. Changed in translation: mRNA recoding by -1 programmed ribosomal frameshifting. Trends Biochem Sci. 2015;40:265–74.

[13] Gesteland RF, Atkins JF. Recoding: dynamic reprogramming of translation. Annu Rev Biochem. 1996;65:741–68.

[14] Mejlhede N, Atkins JF, Neuhard J. Ribosomal -1 frameshifting during decoding of Bacillus subtilis cdd occurs at the sequence CGA AAG. J Bacteriol. 1999;181:2930–7.

[15] Chen H, Bjerknes M, Kumar R, Jay E. Determination of the optimal aligned spacing between the Shine-Dalgarno sequence and the translation initiation codon of Escherichia coli mRNAs. Nucleic Acids Res. 1994;22:4953–7.

[16] Devaraj A, Fredrick K. Short spacing between the Shine-Dalgarno sequence and P codon destabilizes codon-anticodon pairing in the P site to promote +1 programmed frameshifting. Mol Microbiol. 2010;78:1500–9.

[17] Zavialov AV, Hauryliuk VV, Ehrenberg M. Splitting of the posttermination ribosome into subunits by the concerted action of RRF and EF-G. Mol Cell. 2005;18:675–86.

[18] Fredrick K, Noller HF. Accurate translocation of mRNA by the ribosome requires a peptidyl group or its analog on the tRNA moving into the 30S P site. Mol Cell. 2002;9:1125–31.

[19] Fredrick K, Noller HF. Catalysis of ribosomal translocation by sparsomycin. Science. 2003;300:1159–62.

[20] McGarry KG, Walker SE, Wang H, Fredrick K. Destabilization of the P site codon-anticodon helix results from movement of tRNA into the P/E hybrid state within the ribosome. Mol Cell. 2005;20:613–22.

[21] Spiegel PC, Ermolenko DN, Noller HF. Elongation factor G stabilizes the hybrid-state conformation of the 70S ribosome. RNA. 2007;13:1473–82.

[22] Frank J, Gonzalez RL, Jr. Structure and dynamics of a processive Brownian motor: the translating ribosome. Annu Rev Biochem. 2010;79:381–412.

[23] Ling C, Ermolenko DN. Structural insights into ribosome translocation. Wiley Interdiscip Rev RNA. 2016;7:620–36.

[24] Ermolenko DN, Majumdar ZK, Hickerson RP, Spiegel PC, Clegg RM, Noller HF. Observation of Intersubunit Movement of the Ribosome in Solution Using FRET. J Mol Biol. 2007;370:530–40.

[25] Cornish PV, Ermolenko DN, Noller HF, Ha T. Spontaneous intersubunit rotation in single ribosomes. Mol Cell. 2008;30:578–88.

[26] Ermolenko DN, Noller HF. mRNA translocation occurs during the second step of ribosomal intersubunit rotation. Nat Struct Mol Biol. 2011;18:457–62.

[27] Ling C, Ermolenko DN. Initiation factor 2 stabilizes the ribosome in a semirotated conformation. Proc Natl Acad Sci U S A. 2015;112:15874–9.

[28] Belardinelli R, Sharma H, Caliskan N, Cunha CE, Peske F, Wintermeyer W, et al. Choreography of molecular movements during ribosome progression along mRNA. Nat Struct Mol Biol. 2016;23:342–8.

[29] Peske F, Savelsbergh A, Katunin VI, Rodnina MV, Wintermeyer W. Conformational changes of the small ribosomal subunit during elongation factor G-dependent tRNA-mRNA translocation. J Mol Biol. 2004;343:1183–94.

[30] Walker SE, Shoji S, Pan D, Cooperman BS, Fredrick K. Role of hybrid tRNA-binding states in ribosomal translocation. Proc Natl Acad Sci U S A. 2008;105:9192–7.

[31] Shi X, Chiu K, Ghosh S, Joseph S. Bases in 16S rRNA important for subunit association, tRNA binding, and translocation. Biochemistry. 2009;48:6772–82.

[32] Studer SM, Feinberg JS, Joseph S. Rapid kinetic analysis of EF-G-dependent mRNA translocation in the ribosome. J Mol Biol. 2003;327:369–81.

[33] Salsi E, Farah E, Ermolenko DN. EF-G Activation by Phosphate Analogs. J Mol Biol. 2016;428:2248–58.

[34] Svidritskiy E, Ling C, Ermolenko DN, Korostelev AA. Blasticidin S inhibits translation by trapping deformed tRNA on the ribosome. Proceedings of the National Academy of Sciences of the United States of America. 2013;110:12283–8.

[35] Korostelev A, Trakhanov S, Asahara H, Laurberg M, Lancaster L, Noller HF. Interactions and dynamics of the Shine Dalgarno helix in the 70S ribosome. Proc Natl Acad Sci U S A. 2007;104:16840–3.

[36] Yusupova G, Jenner L, Rees B, Moras D, Yusupov M. Structural basis for messenger RNA movement on the ribosome. Nature. 2006;444:391–4.

[37] Noller HF, Lancaster L, Zhou J, Mohan S. The ribosome moves: RNA mechanics and translocation. Nat Struct Mol Biol. 2017;24:1021–7.

[38] Tenson T, Hauryliuk V. Does the ribosome have initiation and elongation modes of translation? Mol Microbiol. 2009;72:1310–5.

[39] Verma M, Choi J, Cottrell KA, Lavagnino Z, Thomas EN, Pavlovic-Djuranovic S, et al. A short translational ramp determines the efficiency of protein synthesis. Nat Commun. 2019;10:5774.

